# Fitness Landscape of the Fission Yeast Genome

**DOI:** 10.1101/398024

**Authors:** Leanne Grech, Daniel Charlton Jeffares, Christoph Yves Sadée, María Rodríguez-López, Danny Asher Bitton, Mimoza Hoti, Carolina Biagosch, Dimitra Aravani, Maarten Speekenbrink, Christopher J. R. Illingworth, Philipp H. Schiffer, Alison L. Pidoux, Pin Tong, Victor A. Tallada, Robin Allshire, Henry L. Levin, Jürg Bähler

**Affiliations:** Department of Genetics, Evolution and Environment, Gower Street – Darwin Building, University College London, London, WC1E 6BT, UK; Department of Biology, University of York, Wentworth Way, York, YO10 5DD, UK; Centro Andaluz de Biología del Desarrollo, Universidad Pablo de Olavide/Consejo Superior de Investigaciones Científicas, Carretera de Utrera Km1, 41013, Seville, Spain; Experimental Psychology, University College London, 26 Bedford Way, London, WC1H 0AP, UK; Department of Genetics, University of Cambridge, Downing Street, Cambridge, CB2 1EH, UK; Wellcome Trust Centre for Cell Biology, University of Edinburgh, Michael Swann Building, Max Born Crescent, Edinburgh, EH9 3BF, UK; Division of Molecular and Cellular Biology, Eunice Kennedy Shriver National Institute of Child Health and Human Development, National Institutes of Health, Bethesda, MD 20892, USA; UCL Genetics Institute, University College London, London, WC1E 6BT, UK

**Keywords:** *Schizosaccharomyces pombe*, fission yeast, transposon mutagenesis, TraDIS, Tn-Seq, fitness landscape

## Abstract

**Background:** Non-protein-coding regions of eukaryotic genomes remain poorly understood. Diversity studies, comparative genomics and biochemical outputs of genomic sites can be indicators of functional elements, but none produce fine-scale genome-wide descriptions of all functional elements.

**Results:** Towards the generation of a comprehensive description of functional elements in the haploid *Schizosaccharomyces pombe* genome, we generated transposon mutagenesis libraries to a density of one insertion per 13 nucleotides of the genome. We applied a five-state hidden Markov model (HMM) to characterise insertion-depleted regions at nucleotide-level resolution. HMM-defined functional constraint was consistent with genetic diversity, comparative genomics, gene-expression data and genome annotation.

**Conclusions:** We infer that transposon insertions lead to fitness consequences in 90% of the genome, including 80% of the non-protein-coding regions, reflecting the presence of numerous non-coding elements in this compact genome that have functional roles. Display of this data in genome browsers provides fine-scale views of structure-function relationships within specific genes.

## Background

A goal of genetics is to understand what sequence elements within genomes specify cellular and organismal function. The highly-transcribed protein-coding regions of eukaryote genomes are routinely detected within genomes and are well studied. The numerous non-coding elements, on the other hand, are more challenging to detect, profile and functionally describe. While biochemical assays of genome activity can indicate functional units, inferring function based *solely* on biochemical activity, e.g. the ENCODE project’s definition of functional DNA [1], is inconsistent with evolutionary analysis that show no signal of conservation for substantial proportions of larger eukaryotic genomes [2,3].

In theory, functionally important elements could be detected by their conservation between lineages relative to neutral elements. However, such analyses suffer from the paradox that more divergent species allow more sensitive detection of small functional elements, but there will be fewer shared functional regions [4]. Similarly, patterns of diversity detect evolutionarily constrained regions within a species [5-7]. However, these analyses are limited to summaries of annotation types, rather than defining particular conserved elements, because segregating genetic variants are generally too sparse within specific genes to estimate the fitness effects of mutations accurately. Additionally, various factors can affect segregating variants and/or allele frequencies at any particular genomic locus, including recombination rate [8] and recent events of selection which purge diversity in surrounding areas [9,10]. For these reasons, neither diversity nor divergence analyses have sufficient power to describe functional constraint at gene or sub-genic resolution. In contrast, high-density transposon-insertion libraries generated from independent repeats can precisely define functional elements and have provided estimators of gene-knockout fitness in bacterial genomes [11-15].

To define functional elements in a eukaryote genome, we generated multiple dense insertion libraries in fission yeast (*Schizosaccharomyces pombe*), using the *Hermes* cut and paste transposon system [16]. We developed a HMM to account for biases in insertion frequency and smooth the stochastic insertion profiles into meaningful measures of insertion-fitness profiles that span multiple continuous genome positions. We analysed this data with respect to genome annotation, genetic diversity, divergence and transcriptional output. This study provides a detailed resource for the understanding and analysis of non-genic functional regions in this model species. This analysis shows that even this well-annotated genome features abundant non-coding functional elements that have not previously been recognized. It provides a detailed resource for further study of genic and non-genic functional elements.

## Results

### Generation of Dense *Hermes* Insertion Libraries in Fission Yeast

We generated nine *Hermes* insertion libraries using modifications of previously published methods [16-18]. Insertions were generated in cultures undergoing rapid mitotic proliferation, serially diluted for approximately 25 generations **(supplementary fig. 1)**. Insertion sites were identified using a custom *Hermes*-end primed sequencing strategy to produce paired-end reads **(supplementary fig. 2)**. This approach included the attachment of a 10-nucleotide (nt) unique molecular identifier (UMI) to each sequenced DNA molecule, which enabled us to remove PCR-generated duplicates of *Hermes*-containing DNA molecules and thus count the number of insertions per position. These counts represent either multiple independent insertions at a genomic location (in different cells within a library), or the result of a single insertion event that has been propagated by cell division.

The libraries contained an average of 1.8 million genomic insertions **(supplementary table 1)**. Collectively, our libraries contained 31 million insertions at 930,000 unique sites, an average insertion density of 1 insertion site per 13 nt of the genome.

### Insertion Density is Consistent with Expectations of Functional Constraint

Based on previous transposon analyses in bacteria and yeasts, we expected that more important regions would tolerate fewer insertions [14,18,19]. Initial analysis showed that both insertion density (unique insertion positions/site) and average insertion count (insertion instances per site) were significantly lower in essential genes compared to non-essential genes and higher in non-genic regions **(supplementary fig. 3)**. This result suggested that insertions reflect the relative functional importance of these annotated elements.

Notably, the mitochondrial genome also featured high insertion density, but with little difference between coding and non-coding regions **(supplementary fig. 4)**. This result likely reflects that any given transposon insertion among multiple mitochondrial genomes will have little or no consequence for the cell. Nevertheless, this finding shows that *Hermes* transposition can readily occur in mitochondria.

To systematically examine the relationship between genomic regions and insertions, we compared our *Hermes* insertion data with genetic diversity (π), both within the species and divergence between *Schizosaccharomyces* species. Based on these evolutionary measures of functional constraint, we divided the genome into four annotation classes: coding regions of essential genes, coding regions of non-essential genes, 5’/3’-untranslated regions (UTRs) and introns, and genomic regions with no annotation (generally intergenic regions). The relative levels of genetic diversity and divergence consistently showed that essential coding regions were subject to higher constraint than non-essential coding regions, followed by UTRs/introns, with unannotated regions being the least constrained. *Hermes* insertion density (unique insertion positions/100 nt) and mean insertion count were consistent with this ranking **(fig. 1)**. These findings indicate that analysis of *Hermes* insertions can quantify the fitness profiles of both coding and non-coding regions.

**Fig. 1.**
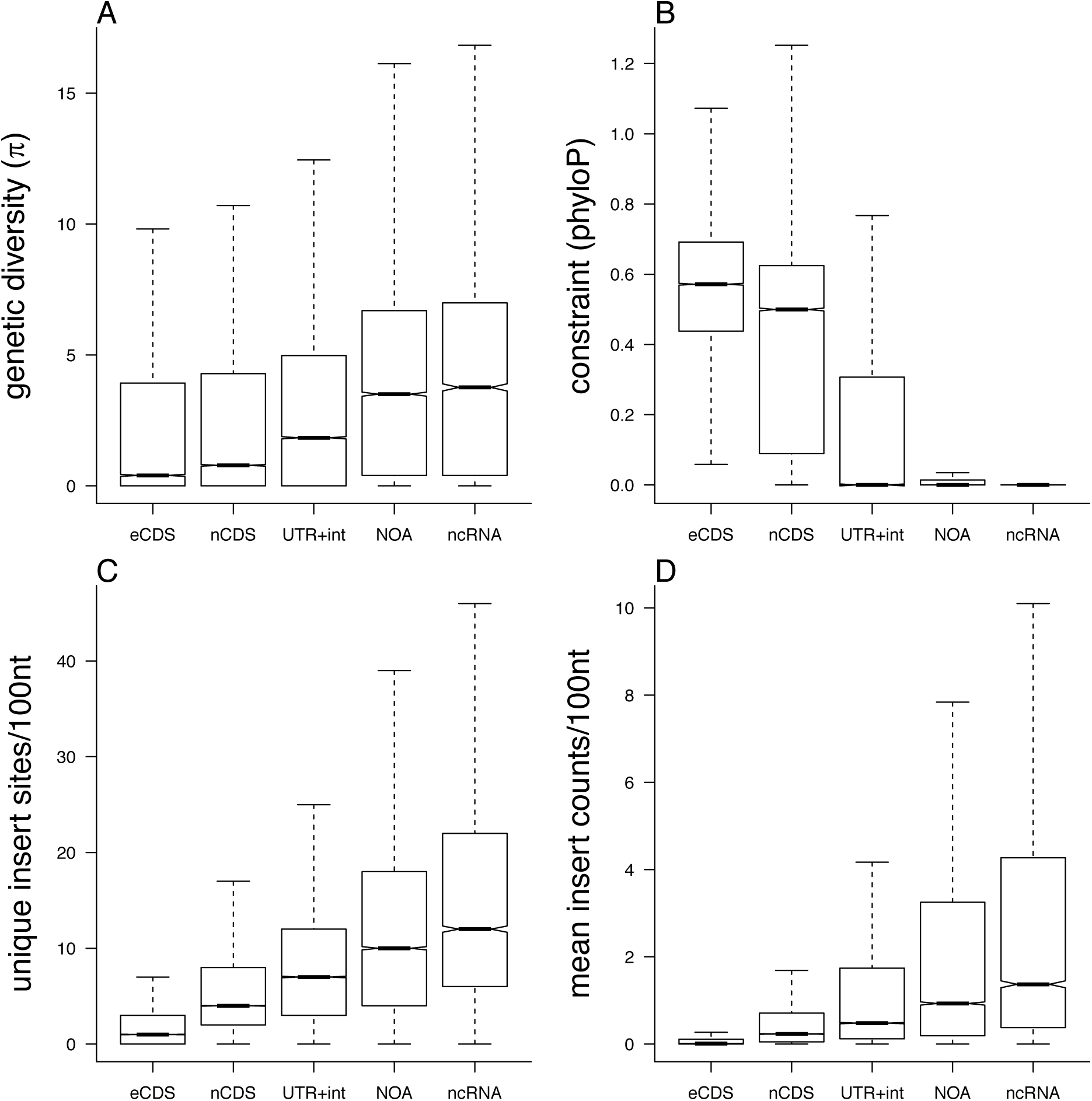
*Hermes* insertion data recapitulate signals of evolutionary constraint. For protein-coding regions of essential genes (eCDS), protein-coding regions of non-essential genes (nCDS), 5’/3’ UTRs and introns (UTR+int), regions of the genome without any annotation (NOA) and non-coding RNAs ncRNAs) we show: **(A)** the genetic diversity from 57 strains of *S. pombe* [5], measured in 100 nt windows, and **(B)** the phyloP measure of constraint [20] between four *Schizosaccharomyces* species (mean phyloP score, over 100 nt windows). Similarly, for pooled proliferation *Hermes* data, we show: **(C)** the number of unique insertion sites/100 nt, and **(D)** the mean insertion counts/100 nt (calculated including sites without insertions as zero counts).

### Application of a Hidden Markov Model to Account for Insertion Biases

Previous analyses have shown that the *Hermes* transposon insertions are biased towards nucleosome-free DNA and that they preferentially occur in DNA with a degenerate sequence motif (TNNNNA) [18,21]. We sought to develop a prediction of the fitness consequences of transposon insertions at a fine-scale resolution correcting for such bias. This prediction should also reflect that neighbouring nucleotides in a genome do not function independently but as ‘functional’ units (e.g. exons, introns, UTRs). We developed a HMM to correct for these insertion biases and smooth the signal from stochastic insertions into contiguous functional units. In this model, the observed data are the insertion counts and the ‘hidden’ state is the degree of biological importance. Regions with greater importance are expected to have fewer insertions.

Our model utilised measurements of nucleosome density and sequence composition. Genome-wide profiles of nucleosome density were obtained from proliferating cells [22]. Next, the sequence composition of previously recorded *in vitro* insertion sites [18] were evaluated to find a degenerate insertion motif. We then constructed a sequence composition measure, termed insertion motif similarity score (IMSS), which describes the similarity of each position in the genome to this motif. Data from these two measurements was used to construct generalised linear models describing the relationship between insertion density, nucleosome density and IMSS **(supplementary fig. 5)**.

Our HMM divided the genome into five states, from state 1 (S1), indicating the sites at which transposon insertion had the greatest negative functional consequences, to state 5 (S5), indicating sites at which insertion had the least negative (or potentially positive) functional consequences. This number of states was obtained from initial trials with the model, detailed below. Annotated regions of the genome were used to train the model. The first state, S1, was trained on coding regions of essential genes (whose knockouts are inviable), S2 was trained on coding regions of non-essential genes, S3 on regions that may have some importance but weaker signals (introns and UTRs), S4 on unannotated intergenic regions that show high genetic diversity [5], where mutations or insertions may be neutral, and S5 on the top-10% insertion-dense sites to allow for the possibility that insertions in some positions enhance cell survival.

The model was fitted to the data by maximum likelihood, using the EM algorithm. The Viterbi algorithm was then used to determine the most likely state (S1-S5) for each genomic position given the nucleosome density, IMSS, and insertion counts. Model fitting did not explicitly include annotations (see Methods for details on HMM). HMM states were highly consistent between independent HMM model fitting runs (see Methods). Insertion data, HMM states, nucleosome density and conservation measures are available in a dedicated genome browser http://bahlerweb.cs.ucl.ac.uk/bioda and in the fission yeast model organism database PomBase (www.pombase.org). These tools allow users to check functional information for regions of interest, including fine-scale structure-function relationships within specific genes and putative regulatory regions.

### Fitness Consequences of Insertions

Transposon insertions had negative fitness consequences over most of the genome, with 91% of the genome being assigned to states S1 or S2. Protein-coding regions of essential genes, used as training data for S1 sites, feature both high between-species conservation and low within-species diversity **(fig. 1)**. The HMM assigned 87% of these regions to S1 **(fig. 2)**, along with 32% of non-essential protein-coding regions.

**Fig. 2.**
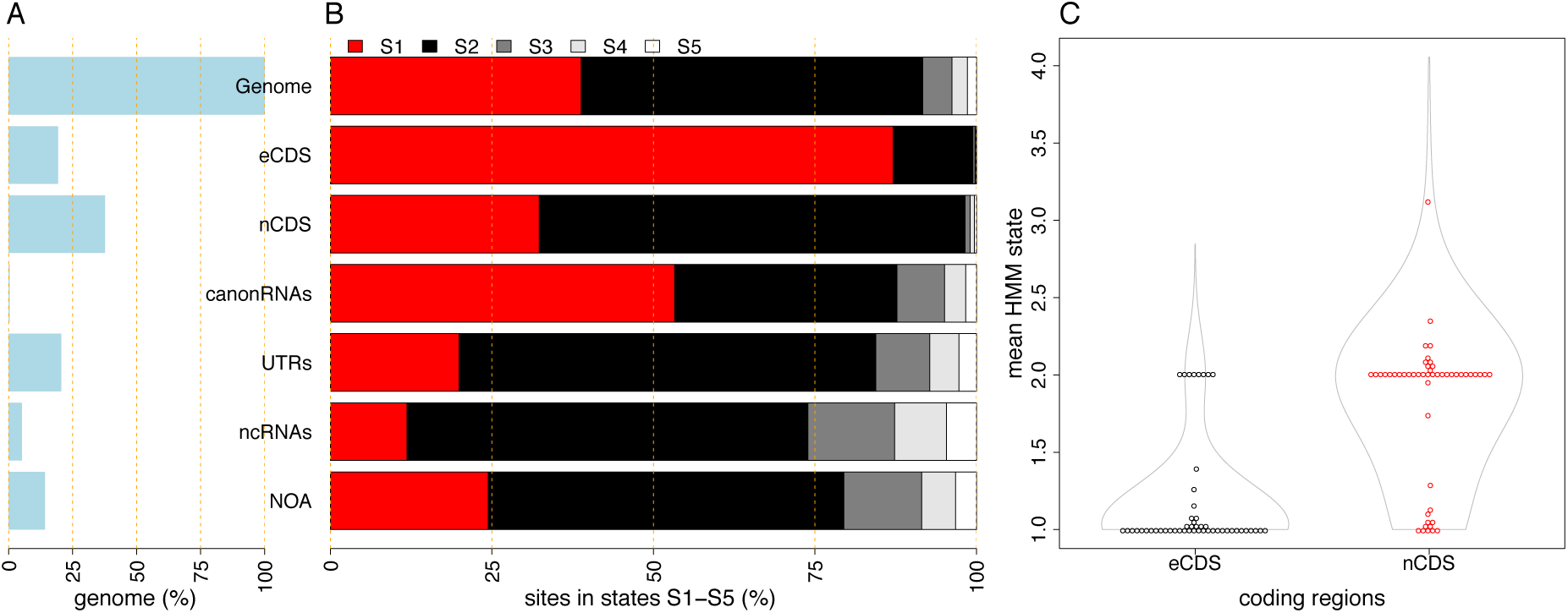
Functional Landscape by Annotation Type. The HMM defined five states based on *Hermes* transposon insertions. State 1 (S1) refers to the most important regions, with the least insertions, and state 5 (S5) with the highest density of insertions. **(A)** Percentage of *S. pombe* genome covered by various annotation types: entire genome (100%), essential protein-coding regions (eCDS), protein-coding non-essential regions (nCDS),canonical non-coding RNAs (snRNAs, snpRNAs, tRNAs, rRNAs, canonRNAs), 5’/3’-UTRs (UTRs), non-coding RNAs (ncRNAs), and unannotated regions (no-anno). **(B)** Proportions of each annotation type in the five states: S1 (red), S2 (black), S3 (dark grey), S4 (light grey) and S5 (white). **(C)** Mean HMM states for essential (eCDS) and non-essential (nCDS) coding regions. Representative 50 points are shown for each type to indicate that most essential coding regions have mean state ∼1 (85% mean state <1.2).

Our analysis indicates that most of the non-coding genome in this species encodes functional elements. The fission yeast genome is much more compact compared to mammalian and plant genomes, with 42% of the current annotation not coding for proteins or canonical non-coding RNAs (ncRNAs); including 20% UTRs, 5% other ncRNAs that do not overlap and protein-coding genes, and 14% with no functional annotation at present. New analysis has discovered almost 6000 new ncRNAs [23], indicating that many functional units remain undescribed.

The HMM assigned 82% non-protein-coding regions to S1 or S2, indicating that they were strongly insertion-depleted relative to genome-wide expectations. UTRs, ncRNAs and unannotated regions were each also insertion-depleted to some extent. **(fig. 2A, B)**. This measure far exceeds the proportion that would be defined as important with the limited comparative genomics data available. For example, 24% of regions with no functional annotation are strongly insertion-depleted (S1), yet these regions show very little conservation between *Schizosaccharomyces* species **(fig. 1).** We also observe that ∼12% of the positions within essential genes contain sufficient insertions to be assigned HMM state 2. These regions could be a mix of two components: annotation mistakes, or could reflect non-essential domains within essential proteins, as described in budding yeast [19].

### HMM states predict the fitness costs of protein-coding gene disruption

To examine whether the HMM contained information about the relative fitness cost of gene disruption, we calculated the mean HMM state for each protein-coding gene. While essential coding genes had much lower mean states **(fig. 2C)**, essential and non-essential genes showed overlapping distributions. To assess the validity of this measure, we compared it to the colony sizes of viable knockout mutants on solid media, an orthogonal measure of gene disruption fitness alteration that uses different media, a more direct fitness measure, and different methods to obtain complete gene deletions [24]. Reassuringly, the mean HMM state positively correlated with the colony size of knockout mutants (Pearson *r* = 0.34, P = 10^−90^, **fig. 3A**) [25,26]. Genes with fewer insertions (lower mean HMM states) were also more likely to be conserved between *Schizosaccharomyces* species and highly expressed (**fig. 3B, C)**, both expectations for genes that cause strong fitness consequences when mutated. In summary, these analyses show that the insertion and analysis methods recover biologically meaningful fitness measures that add value beyond the binary classification of essential/non-essential genes that can be obtained from whole-gene disruptions.

**Figure 3.**
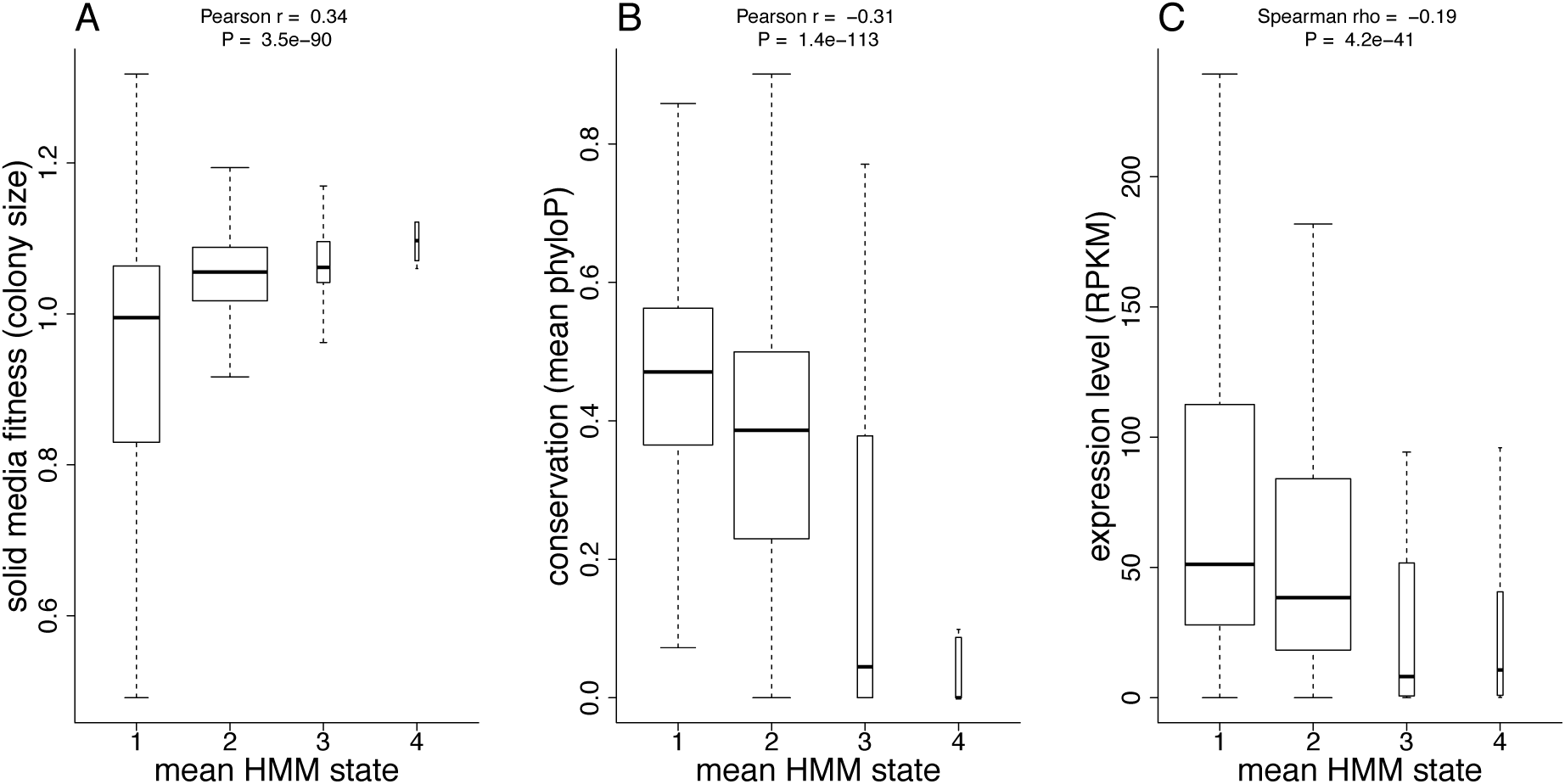
Gene mean HMM states are estimators of gene disruption fitness. Protein-coding genes classified into four categories by the mean HMM states, showing those that are ∼1 (< 1.5), ∼2 (> 1.5 and ≤ 2.5), ∼3 (>2.5 and ≤ 3.5) and ∼4 (>3.5 and ≤ 4.5). Mean HMM states were positively correlated with solid media fitness **(A)**, an orthogonal measure. Mean HMM states were also negatively correlated with conservation (lower HMM states were more conserved) **(B)**, and negatively correlated with gene expression (lower HMM states were more highly expressed) **(C)**.

### HMM-Defined Functional Elements

To examine whether the HMM states captured previously annotated elements, such as introns, promoters, and protein-coding exons, we defined 256,815 ‘HMM-defined elements’ (HDEs) as genomic regions that feature a continuous run of one HMM state. All S4 or S5 HDEs were less than 100 nt, and mostly intergenic, indicating that only short regions in this genome can tolerate insertions without affecting fitness.

We excluded these S4/S5 HDEs from further analysis, leaving 10,015 HDEs with a median length of 618 nt, which account for 90% of the mappable genome. HDE edges were closer to edges of existing annotations than expected by chance (Wilcoxon Rank Sum test, P <10^−16^, **fig. 4A, B**). This result is consistent with these HMM-defined regions representing boundaries of a variety of biologically-relevant elements (including transcriptional units, spliced exons, protein-coding regions).

**Fig 4.**
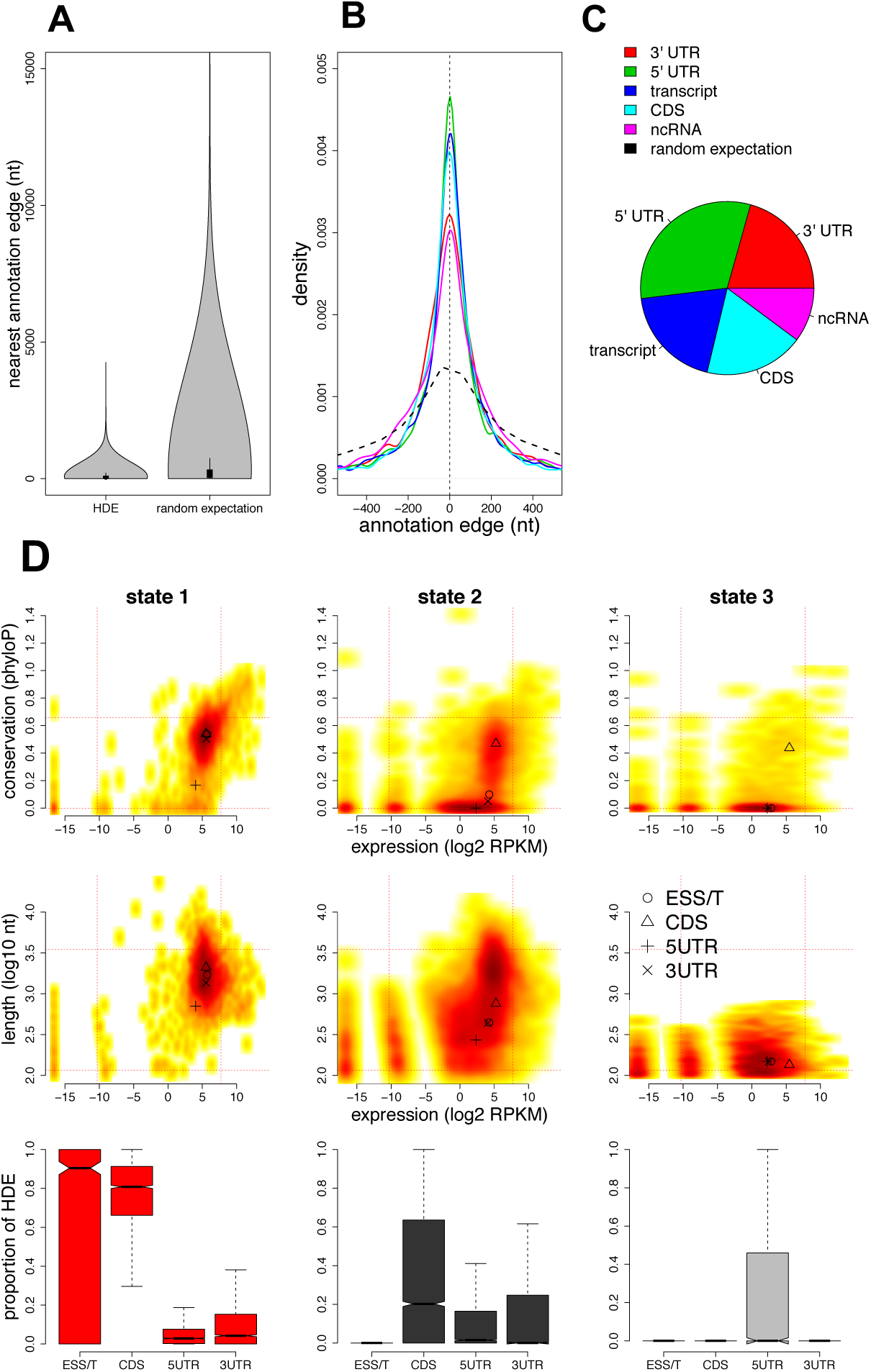
HMM-defined elements describe functional genomic outputs. Parts A-C show that the boundaries of HMM-Defined Elements (HDEs) are aligned to or close to the boundaries of existing annotations, as defined in the legend at top right. The random expectation is derived from the same number of elements of the same lengths, placed at random on the genome. **(A)** HDEs have a smaller distance to the nearest annotation than the random expectation. **(B)** For all HDE edges we show the distance to the nearest annotation type, including 5/3’ UTRs, transcripts (transcription start/stop positions), coding sequences (amino-acid encoding regions, CDS), non-coding RNAS (ncRNAs), with lines coloured according to the legend at right. **C)** HDEs fell closest to a variety of annotations. The pie chart shows the proportions of nearest annotations, indicating a bias towards defining 5’UTR edges. There were subtle differences between S1, S2 and S3 states in this respect (not shown). **(D)** Density plots describe various characteristics of HDEs, from left showing S1, S2 and S3 HDEs. Conservation (*y* axis, top row) levels are mean phyloP measures from four *Schizosaccharomyces* species. HDE lengths (y axis, middle row) are shown on a log_10_ scale. Expression levels (x axes) are RNA-Seq RPKMs from proliferating cells. Dashed horizontal and vertical lines show the 5^th^ and 95^th^ percentiles of conservation, expression levels or lengths. The positions of symbols (circle, triangle *etc.*) indicate the median positions within each state for essential transcripts (ESS/T), coding regions (CDS), and 5’/3’ UTRs. For example, the few conserved S3 sites are coding regions. The bottom row shows the proportion of HDEs that are annotated as essential transcripts (ESS/T), protein-coding sequence (CDS), 5’ UTR and 3’ UTR.

To characterise these HDEs, we calculated their conservation during evolution and their RNA expression levels. The HDEs which were most insertion-depleted, and therefore most critical for cell function (S1 elements), covered 35% of the mappable genome. These HDEs showed distinct features: they were most conserved between species, the longest (mean length 1.9 kb), most highly expressed, and generally composed of protein-coding regions **(fig. 4D)**. Another 52% of the genome was composed of S2 elements (mean length 1.0 kb), including mainly coding regions and UTRs, which also showed relatively high expression levels and conservation. The inclusion of many 5’- and 3’-UTRs in S2 elements indicates that these non-coding regions often contain regulatory sites whose disruption impairs cellular function. Finally, the S3 elements occupied only 3% of the genome, were seldom conserved, generally short (mean length 0.18 kb), and almost exclusively 5’-UTRs. These UTRs likely contain regulatory sites, because they feature fewer insertions than S4 regions, but would have been difficult to identify without the insertion data because they are neither conserved nor very highly transcribed. As the *Schizosaccharomyces* clade contains only four species, subtle constraint will likely remain undetected. Overall, 10% of the important sites in the genome (S1-S3) showed no signal of conservation between species.

## Discussion

Dense transposon-insertion libraries can identify genes whose disruption affects fitness (in particular conditions) within bacterial genomes with high resolution [11-15]. However, similarly high-resolution descriptions of eukaryotic genomes are more limited, and have not yet achieved nucleotide-level definitions of fitness landscapes [18,19]. Studies with eukaryotic genomes are also more challenging, because they are larger and contain nucleosomes, which bias integration rates. With the density of our insertions in libraries from proliferating cells (26.7 million insertions, 1 unique insertion site/13 nt), and the application of a HMM to account for insertion bias, we analysed functional importance at near single-nucleotide resolution.

The findings of the HMM are validated by the demonstration that continuous single-state genome sections (HMM-defined elements, HDEs) are closely aligned to existing annotations, and define elements with different properties (**fig. 4**). As the *Hermes* insertion data recapitulates signals of genetic diversity and divergence within different annotation categories, we can be confident that insertion density reflects functional constraint **(fig. 1)**. The application of a hidden Markov Model robustly accounted for insertions biases, since HMM states strongly depended on insertion density but only weakly correlated with nucleosome density and nucleotide motif **(supplementary fig. 6)**.

Our HMM analysis of transposon insertions assigned 91% of the fission yeast to HMM S1 or S2 (which were trained on essential and non-essential coding regions, respectively). Based on this, we conclude that 91% of the genome contains functional elements that are affected by transposon insertions. These likely functional regions of the genome include 80% of the currently un-annotated genome, consistent with the presence of many unrecognised functional elements in non-coding regions of this model organism. This is the first near nucleotide-level study of fitness consequences in a eukaryote genome, so there are no clear expectations. In theory, species with larger population sizes are expected to maintain smaller genomes with larger proportions of functional DNA [27]. Consistent with this prediction, analysis of comparative genomics data has estimated that 5-15% of the human genome shows signals of conservation [28-30], whereas increasingly larger proportions of the *Drosophila* (∼50%), *Caenorhabditis* (37%), and *Saccharomyces* yeast (up to 68%) genomes are conserved [31]. Our estimate of functional regions is likely larger due to the limitation of comparative genomics, that is it only able to detect regions that have continuously subject to purifying selection throughout the phylogeny of the species aligned [4]. It is also possible that in some cases transposon insertions can disrupt the function of larger neighbouring regions, although the sites of insertions themselves are not functional.

A limitation of our study is that the transposon method does not reveal how non-coding genomes elements function. Future work will reveal whether these elements function as the widespread non-coding transcripts [22] and/or as regulatory elements controlling the expression of coding genes.

## Conclusion

Our analysis indicates that the fission yeast genome is densely packed with functional elements, including many uncharacterised non-protein-coding elements. We estimate that 90% of the genome contains functional elements that are impaired by transposon insertions, including 80% of the non-protein-coding regions. We expect that saturating transposon mutagenesis data has potential to define functional non-protein-coding elements within eukaryote genomes that would be difficult to detect with any other contemporary method.

## Methods

### Creating *Hermes* Insertion Libraries

*Hermes* insertion libraries were constructed as described [16] using the pHL2577 and pHL2578 plasmids, except that the transposition frequency was calculated by dividing the number of colonies on YES 5-FOA+G418 plates by the number of colonies on YES plates. All experiments were performed in an *S. pombe* strain with the genotype *ura4*–D18 *leu1–32 h*^*–*^. Typically, <0.2% of cells in libraries contained genomic *Hermes* insertions, so we expect that most insertion mutants contain a single insertion.

### Generating DNA Libraries for Sequencing

Genomic DNA was extracted from insertion libraries using phenol/chloroform extraction. All DNA extracted from a library was processed. DNA was sheared to an average size of 200 bp using a Covaris S2 ultrasonicator (Covaris, Woburn, Massachusetts). Sheared DNA was end repaired using the NEBNext^®^ End Repair Module (NEB, Hitchin, UK). Linker1-Random10mer and Linker2 **(supplementary table 4)** were ligated using the NEBNext^®^ Quick Ligation Module (NEB, Hitchin, UK). In Linker1-Random10mer, the random 10 nt sequence acted as a UMI to distinguish unique chromosomal insertions from PCR amplifications. DNA was then digested with KpnI-HF (NEB, Hitchin, UK) to exclude residual *Hermes* pHL2577 donor plasmid from PCR amplification (as the plasmid contains a unique KpnI site). NEBNext^®^ modules were used according to manufacturer’s instructions. To enrich for fragments containing the *Hermes* transposon, DNA was amplified with BIOTAQ™ DNA polymerase (Bioline, Essex, UK) using a primer that complimentary to the *Hermes* transposon (1-Transposon-4NNNN), and to the linker (Linker1-Amp**, supplementary table 4)**. Ultimately, a second PCR attached the multiplex oligonucleotides for Illumina MiSeq sequencing; the MS-102-2022 MiSeq reagent kit v2 (300 cycles) (Illumina, Cambridge, UK) was used to sequence the libraries. To increase the complexity of the libraries, for each library, ligation and PCR reactions were performed in multiple reactions (in 96-well plates), using a maximum of 1 µg of DNA per well and then re-pooled before sequencing. Detailed protocols are available in the Figshare project *Hermes Transposon Mutagenesis of the Fission Yeast Genome* (will be made publicly available upon manuscript acceptance). Sequence data are available at European Nucleotide Archive in study accession number PRJEB27324. Sample accessions are listed in **supplementary table 5**.

### Computational Processing of Sequencing Data

Bioinformatic processing filtered the sequence data to retain only reads derived from *Hermes* insertions, removed reads with duplicate UMIs, and filtered for correctly-paired high-confidence read-mapping, and ultimately located the positions and orientation (strand) of genomic insertions. Details are as follows. Read 1 architecture was [random4mer][*Hermes*][Genome] (with random 4mer added to increase 5’ Read 1 end complexity to allow Illumina cluster calling). The 4mer was trimmed with fastx_trimmer (http://hannonlab.cshl.edu/fastx_toolkit/). The Reaper tool [32] was used to detect reads with 5’ ends matching the expected *Hermes* sequence, and excluding those within the pHL2577 donor plasmid. Read 2 architecture was [10mer][Linker][Genome]. We used a custom Perl script to exclude duplicate reads with exactly matching 10mers. Processed Reads 1 and 2 were re-paired using Tally [32], and the 10mer and Linker were trimmed with fastx_trimmer. Paired-end reads were aligned to the reference genome [33] and the donor plasmid using BWA-MEM (Li and Durbin 2009). SAMtools [34] was used to select correctly paired reads with a mapping score ≥30 (flags 83 and 99). Finally, we applied custom scripts to identify the location and strand of insertions from the filtered BAM outputs with SAMtools. Insertions found on the same chromosome but on different strands were considered as unique events. Command lines for this procedure and scripts are available in the Figshare project *Hermes Transposon Mutagenesis of the Fission Yeast Genome*, as well as all insertion data, and HMM model fitting results.

### Nucleosome Density Data

The generation of the nucleosome density data has been described in Atkinson et al. [22] and are available at the European Nucleotide Archive under accession number PRJEB21376. The median nucleosome density from two repeats was transformed to a normal distribution. This normalised nucleosome density showed a stronger correlation with insertion density than the raw nucleosome density and was used as a predictor in the HMM.

### Insertion Motif Similarity Score

*In vitro Hermes* insertion data [18] was used to identify a sequence motif corresponding to insertion events in non-nucleosome bound DNA. Strings of 41 nt, centred upon each *in vitro* insertion event were taken from the *S. pombe* reference sequence. The percentage of each nucleotide present at each of the 41 positions was measured and compared to percentage nucleotide compositions calculated across the entire genome. A window of 20 positions was identified for which the composition differed from the genome-wide composition by at least 1% for at least one of the four nucleotides. For each position *i*, we denote the probability of observing the nucleotide *a* as

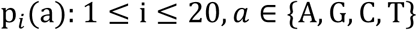

and denote the genome-wide probability of observing the nucleotide *a* as p^gw^(a).

A genome-wide scan was then conducted of strings of 20 consecutive nt in the genome sequence, calculating a likelihood measure of the extent to which each string matched the insertion motif, as compared to the genome-wide base composition. Where a string is given by the nucleotides {a_1_, a_*2*_, …, a_20_} we calculate the insertion motif similarity score as follows:

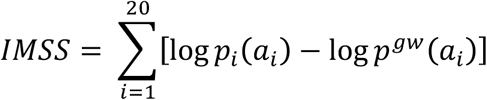

Here a positive score indicates a greater similarity to the insertion motif than to the genome-wide sequence propensity. This likelihood measure was used as a predictor in the HMM.

### Hidden Markov Model

We developed a hidden Markov model using the R package depmixS4 (Visser and Speekenbrink 2010b). These models assume that sequences of observed response variables are dependent on underlying sequences of discrete hidden states. The sequence of hidden states is assumed to follow a first-order Markov process, such that the probability of a state at position *t* depends only on the hidden state at the immediately preceding position *t-*1. The observed responses are assumed conditionally independent given the sequence of hidden states (i.e., correlations between nearby positions are completely accounted for by the hidden states. This model used log_2_-transformed insertion numbers as the observed state. Sites with zero insertions were set to observed state = 0. Each hidden state defined a (zero-inflated) Poisson regression model, with log_2_ insertion count as dependent variable, and the normalised nucleosome density (median of two replicates) and nucleotide preference score as predictors. Missing data for nucleosome density was set to the median. The models parameters (initial state probabilities, state-transition probabilities, and the parameters of the state-dependent zero-inflated Poisson regressions, were estimated by maximum likelihood using the Expectation-Maximisation (EM) algorithm. Initial state distributions were all 1/*n,* where *n* is the number of states. Initial transition matrix was 0.95 for positions remaining in the same state, and 0.05/(n-1) for all other transitions. Initial parameter values of the Poisson regressions were obtained by pretraining each state-dependent model on a subset of the data (see below). These initial parameters were used to start the EM algorithm, the final resulting parameter estimates were determined by maximum likelihood. Neither annotations nor transcriptome data were supplied as predictors to the HMM. Models were fit to the insertion data by the EM algorithm, until convergence of the likelihood (with a tolerance 1×10^−8^) or with a maximum of 150 iterations (since log likelihood fit of models improved little after 150 iterations **(supplementary fig. 7)**.

### Choice of Optimal Model

To select an appropriate number of states and state training data for our HMM, we used ten ‘test data’ subsets of the genome, each a 100 kb fraction as follows: Chromosome I, 100001-200001, 1100001-1200001, 2100001-2200001, 3100001-3200001, Chromosome II, 100001-200001, 1100001-1200001, 2100001-2200001, 3100001-3200001 and Chromosome III, 100001-200001, 1100001-1200001 (test data sets A to J). These regions avoid the chromosome ends, which have unusual properties, such as a high frequency of pseudogenes and native Tf1 transposon insertions [5].

We ran each of the following models on all insertion data from proliferating cells (split into the ten subsets). These models defined the training data in two ways. Firstly, ‘insertion-quantile’ models, where training data was defined solely by the density of unique insertions, calculated over 100 nt windows. For example, a 3-state model split the data into the lower, mid and upper third insertion density for states 1-3. We trialled quantile models from 2 to 10 states. Secondly, annotation-based models. We trialled 2-, 3-, 4-, and 5-state models where the training data was derived from current genome annotations. The 2-state model included coding sequences (S1) and other regions (S2). The 3-state model, coding sequences of essential genes (S1), coding sequences of non-essential genes (S2), introns, unannotated regions, and UTRs (S3). The 4-state model, coding sequences of essential genes (S1), coding sequences of non-essential genes (S2), introns and untranslated regions (S3), and unannotated regions (S4). It differs from the 3-state model in that it differentiates UTRs and introns from unannotated regions. The 5-state model is as the 4-state model, except that it includes a 5th state that contains sites with the highest 10% of unique insertions/100 nt. The response for this state was a Poisson distribution rather than zero-inflated Poisson.

Each of these 13 models was fit (with tolerance 1×10^−8^) to the ten fractions of the genome. Fitting involved optimising the parameter of states at each position, the transition state matrix, and the slope, intercept and zero-fraction of the state model. A 5-state annotation model was chosen as a pragmatic the best fit for running large (million position) data sets. Comparison of the Bayesian information criterion scores (BIC) for 2-5 states indicated that increasing states improved the fit **(supplementary fig. 8)**, but higher state models suffered from increased run times and frequent run failure, and/or highly inconsistent fractions of the subset data assigned to various states (with some states being absent).

Due to the rounding of log_2_ insertion counts, sites with 1 or 0 insertions were set to the same observed state. Rounded log_2_ of insertions+1 (where sites with 0 insertions have different value from those with 1) resulted in a worse fit to the model **(supplementary fig. 9)**.

### Fitting of Chromosome-Wide Data

Once the 5-state annotation model (model 5A) was chosen as a pragmatic best model, it was run on all proliferation libraries, fitting data from five relatively equal portions of the genome separately, to allow runs in a practical time frame and memory. These fractions were: chromosome I left half (positions 1-2789566), chromosome I right half (positions 2789567-5579133), chromosome II left half (positions 1-2269902), chromosome II right half (positions 2269903-4539804), and the entirety of chromosome III (fractions are between 2.26 Mb and 2.79 Mb). The model produced a state prediction for each position in the genome, and the posterior probability of each state at each position. We also fit model 5A to the ageing insertion data (pooled Days 0, 2, 4 and 6) with the same genome subsets.

Collectively, the proliferation samples have a higher count of insertions than any of the pooled ageing libraries (proliferation: 31 million insertions; ageing: 4.6 million insertions). Since training datasets are based on the within-sample insertion densities for each HMM fit, this should account for different densities. Nevertheless, to examine whether this large difference in insertion counts produced radically different fits, we produced a down-sampled dataset from proliferation samples with the same insertions as the ageing sample average (4.5 million insertions). Overall, 85% of sites in this reduced data set were assigned the same state as the full proliferation data, and 98% of sites were within one step of the full data (i.e. full proliferation state +/-1).

These separate fits to the model resulted in similar distributions of states between chromosome arms for both the coding regions and introns of essential genes, supporting consistent convergence of the models between these genome subsets **(supplementary fig. 10, 13)**. To examine whether positions were assigned a consistent state using different subsets of data, and independent fits of the HMM, we made subsets of proliferation (dense data) and ageing Day 6 (less dense data) for the central half of chromosome I (positions 1394783-4184350), which overlaps both the left and right halves used previously. These data were fit to model 5A as before. With dense proliferation data, sites that overlapped the 96.7% of positions were assigned the same state with either left *vs* middle, or right *vs* middle comparisons. For ageing Day 6 data, 97.1% of overlapping positions were assigned the same state. States 1-5 were all consistently assigned (e.g. > 99% of state 5 positions were the same within proliferation data, and similar proportions for all other states). This analysis indicates that these fractions were sufficiently large to preclude fitting to very different local optima. HMM code is available in the Figshare project *Hermes Transposon Mutagenesis of the Fission Yeast Genome*.

### Filtering Badly Mapped Sites

To ensure accurate placement of reads, our pipeline filtered reads mapped with mapping quality ≥30. To avoid the tendency to misinterpret regions that have few insertions due to the loss of low mapping quality, we analysed only sites that had retained ≥90% of the reads (lost <10%) over 500 nt windows after mapping quality filtering. This retained 94.6% of the genome for analysis. After filtering, there was only a weak negative correlation between the HMM state and the proportion of reads filtered (Pearson r =-0.049). All data presented included only the sites that had retained ≥90% of the reads after filtering for Q30 mapping (the ‘mappable genome’).

### Annotation Data

Annotations were from PomBase (ASM294v2, 11/02/2016), including 1538 annotated ncRNAs.

### Transcriptome Analysis

Replicated RNA-Seq data from vegetatively growing, early stationary and deep stationary cultures were retrieved from the European Nucleotide Archive (ENA; http://www.ebi.ac.uk/ena) using the following accession numbers (dataset: PRJEB7403; samples: ERS555567, ERS555607, ERS555570, ERS555612, ERS555571,ERS555613). [22]. Reads were aligned to the *S. pombe* genome as described [35]. The resultant aligned reads were used to compute normalised coverage at the nucleotide level using the genomecov function in the BEDtools suite [36]. Customised R scripts were used to define whether a given region is transcribed.

### Comparative Genomics

We used updated genome assemblies of fission yeasts *S. octosporus, S. japonicus*, and *S. cryophilus* [37]. To improve previous full genome alignments of fission yeast species [38], we incorporated these newly assembled genomes into an alignment with the S. *pombe* genome using progressive-cactus [39] (github version May 2016), using a guide tree based on Rhind *et. al.* [38]. We then applied the phyloP algorithm [40] as implemented in the HAL toolkit [41] to detect constraints. We trained a neutral model using the four-fold degenerate sites from coding regions from the high-quality *S. pombe* annotation.

## Declarations

<u>Ethics approval and consent to participate:</u> N/A

<u>Consent for publication:</u> N/A

<u>Availability of data and material</u>

Sequence data are available at European Nucleotide Archive in study accession number PRJEB27324. Sample accessions are listed in **supplementary table 5**.

Transposon insertion data, R code for the HMM, all other data used for analysis, and detailed protocols are available in the figshare project *Hermes Transposon Mutagenesis of the Fission Yeast Genome* (will be made public after manuscript acceptance).

## Author Contributions

LG produced *Hermes* insertion data, assisted with bioinformatics, statistical analyses and writing the manuscript. DCJ initiated the project, supervised students (LG, CYS, CB, DA), conducted bioinformatics and statistical analyses and wrote the manuscript. CYS assisted with analysis of *Hermes* insertion data. CB and DA produced initial *Hermes* insertion data. DAB implemented the genome browser and analysed RNA-Seq data. MRL produced the nucleosome density data and assisted with production of *Hermes* insertion data. VAT assisted with production of *Hermes* insertion data. MS developed R code for HMM and assisted with statistical analyses. CJRI defined the nucleotide insertion model. PHS aligned and produced conservation measure of *Schizosaccharomyces* genomes. ALP and PT produced assemblies of *Schizosaccharomyces* genomes. RA provided *Schizosaccharomyces* genomes. HLL provided additional *Hermes* insertion data. JB funded the project, supervised the PhD student and postdocs in his group, and helped with writing the manuscript.

## Acknowledgments

We thank Dr Rachel Brown for guidance with *Hermes* methods.

## Supplementary Figures

**Supplementary fig. 1.**
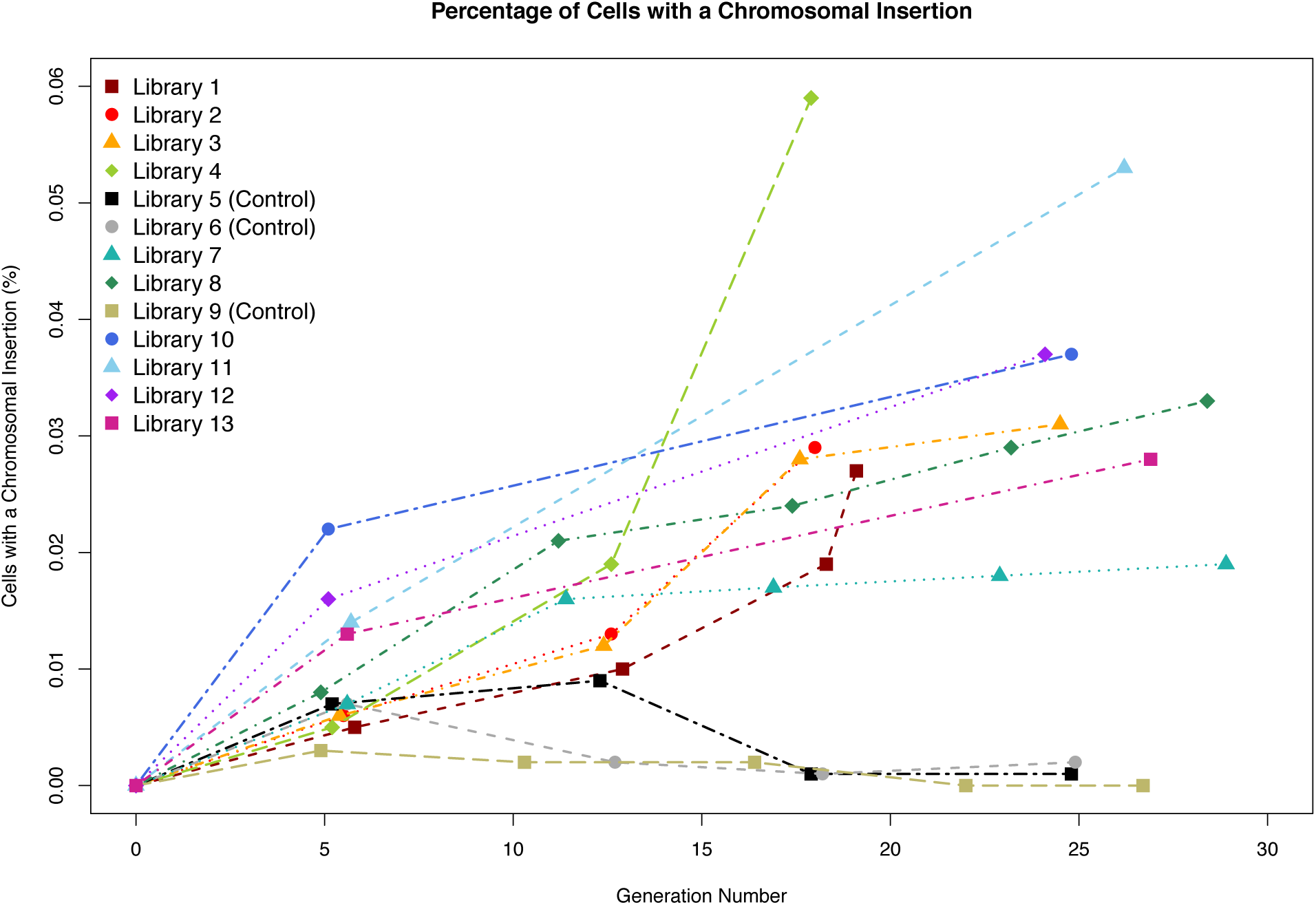
Percentage of cells with a chromosomal insertion. For the nine libraries we generated (and others not described here), we show the percentage of cells with a chromosomal insertion. The proportion was calculated as the number of colonies present on YES + FOA + G418 plates (chromosomal insertions), divided by the number of colonies present on YES plates (all cells).

**Supplementary fig. 2.**
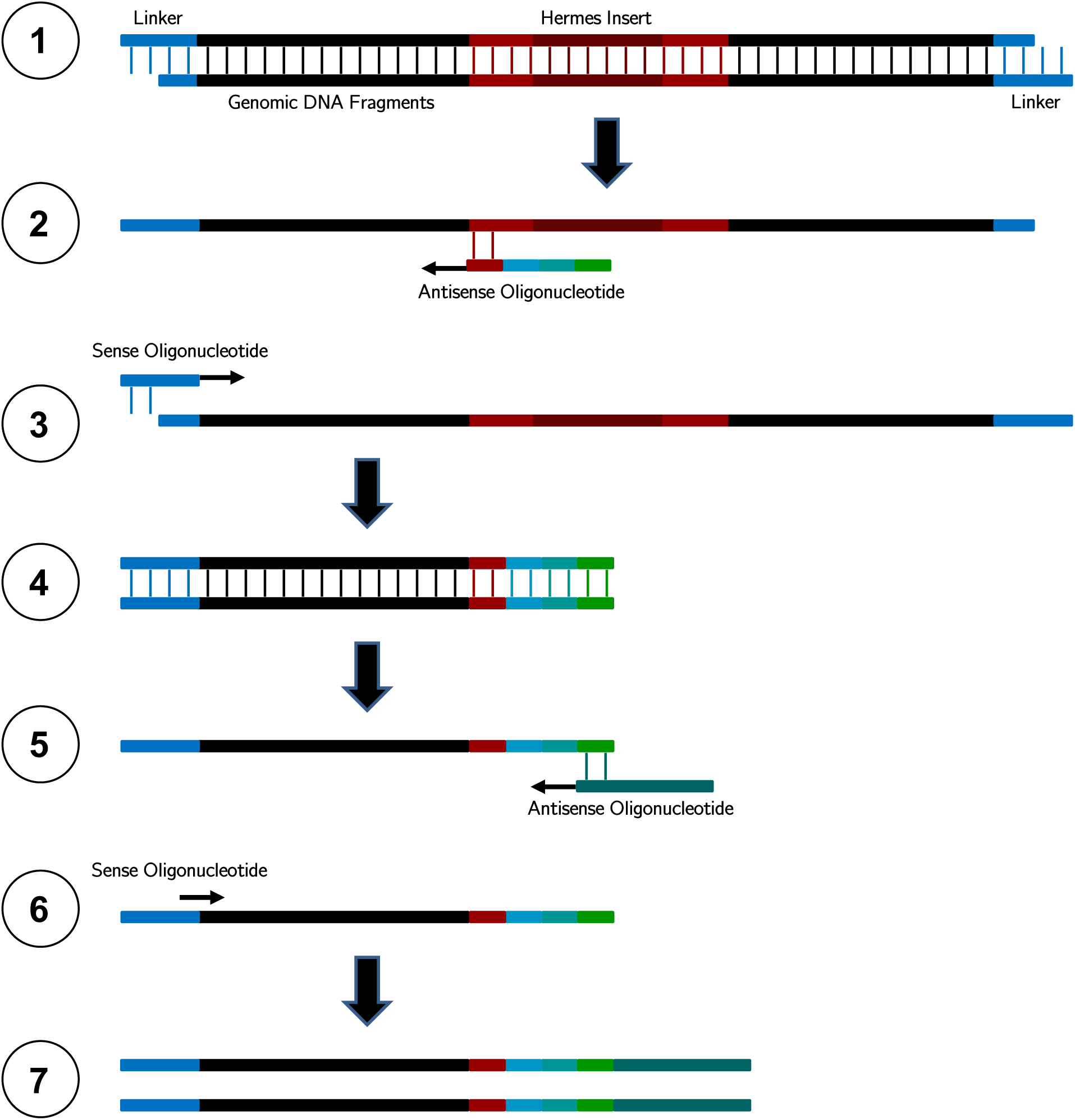
The custom *Hermes*-end primed sequencing strategy. Shows the end-priming strategy used to sequence Hermes-containing fragments. Initially, genomic DNA is extracted, sheared, end repaired, and linkers (Linker1-Random10mer and Linker2) ligated at both terminal ends (1). To enrich for fragments containing the *Hermes* transposon, DNA was amplified with using a primer that is complimentary to the *Hermes* transposon (1-Transposon-4NNNN) (2), and to the linker (Linker1-Amp) (3), to produce fragments that contain linkers, genomic DNA and the *Hermes* right terminal inverted repeat (4). A second PCR attached the multiplex oligonucleotides for Illumina sequencing (5,6), producing the final product that is sequenced (7). Detailed protocols are available in the Figshare project *Hermes Transposon Mutagenesis of the Fission Yeast Genome*.

**Supplementary fig. 3.**
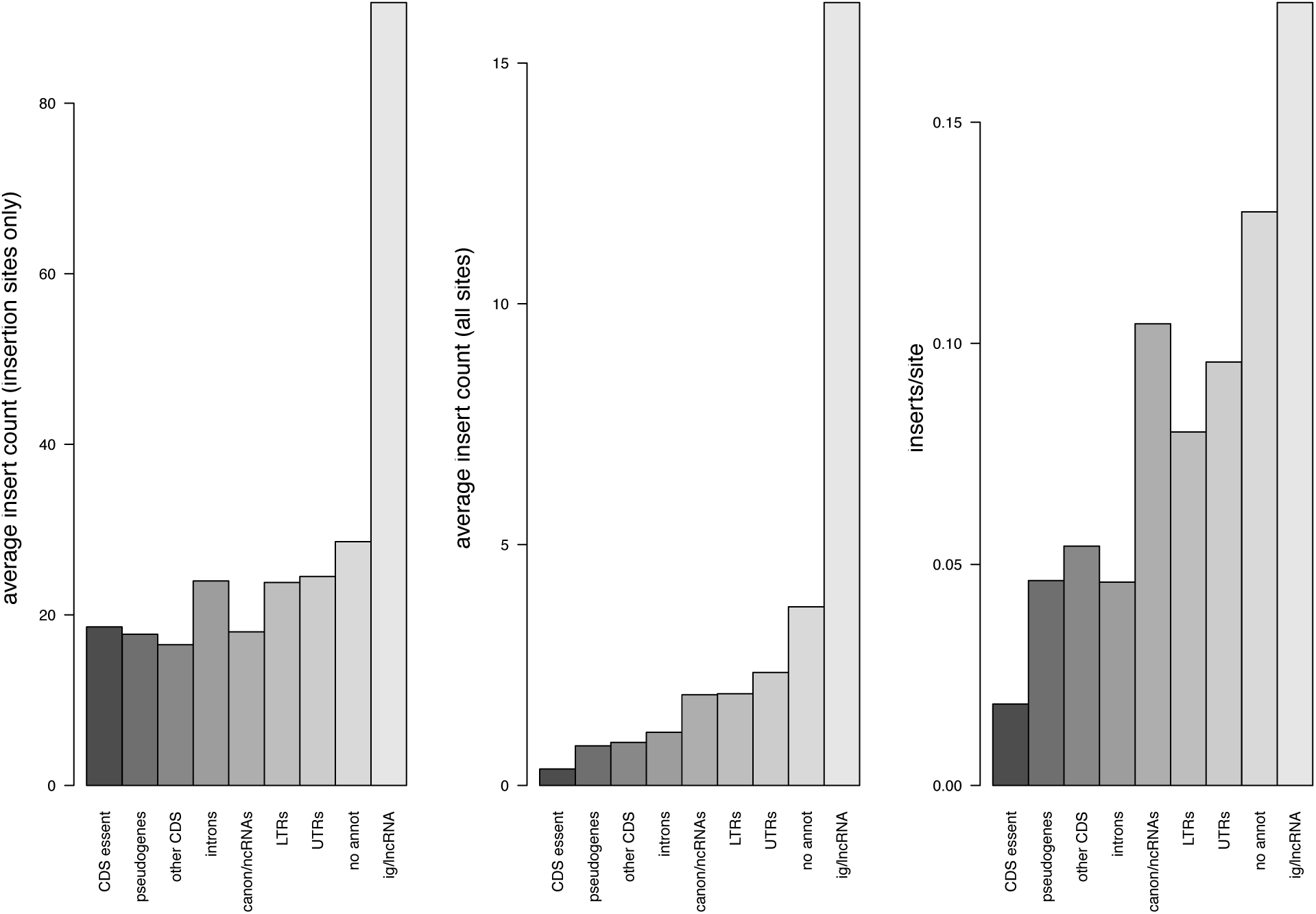
Properties of insertions in different annotation regions. Left panel shows average insertion count in coding regions of essential genes, pseudogenes, other (non-essential) coding regions, introns, canonical non-coding RNAs (snoRNas, tRNAs, rRNAs, snRNAs), long terminal repeats of transposons,5’ and 3’ untranslated regions, regions with no annotation and intergenic long non-coding RNAs. Middle panel shows and average insertion count (all sites, including sites with no insertions) for the same annotation classes. Right panel shows average insertion density (unique insertion positions/site) for the same annotations.

**Supplementary fig. 4.**
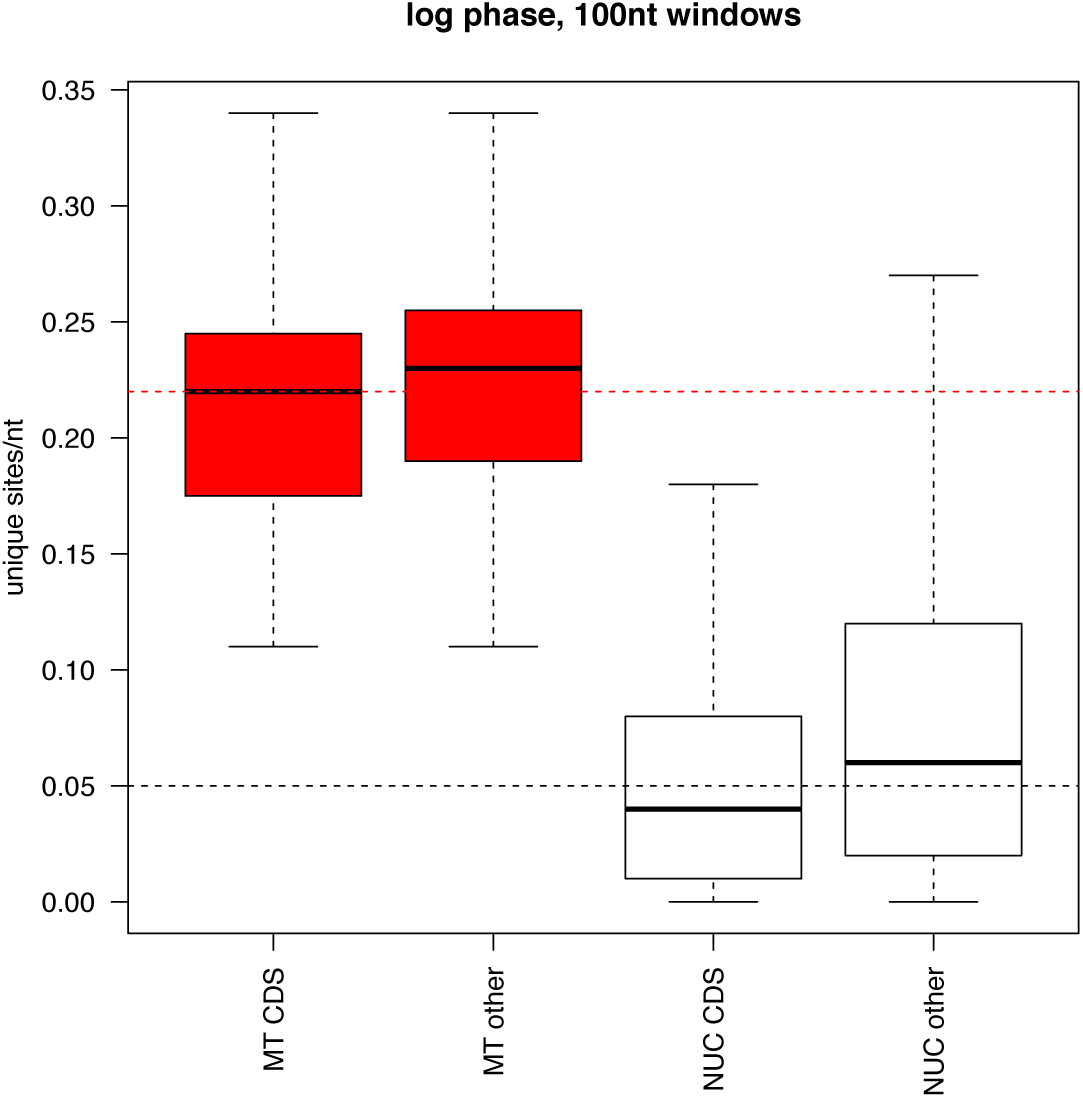
Insertions in the mitochondrial genome. Unique insertions per site in the mitochondrial genome showed little difference between coding and non-coding regions, whereas the nuclear genome showed far fewer insertions in the coding regions.

**Supplementary fig. 5.**
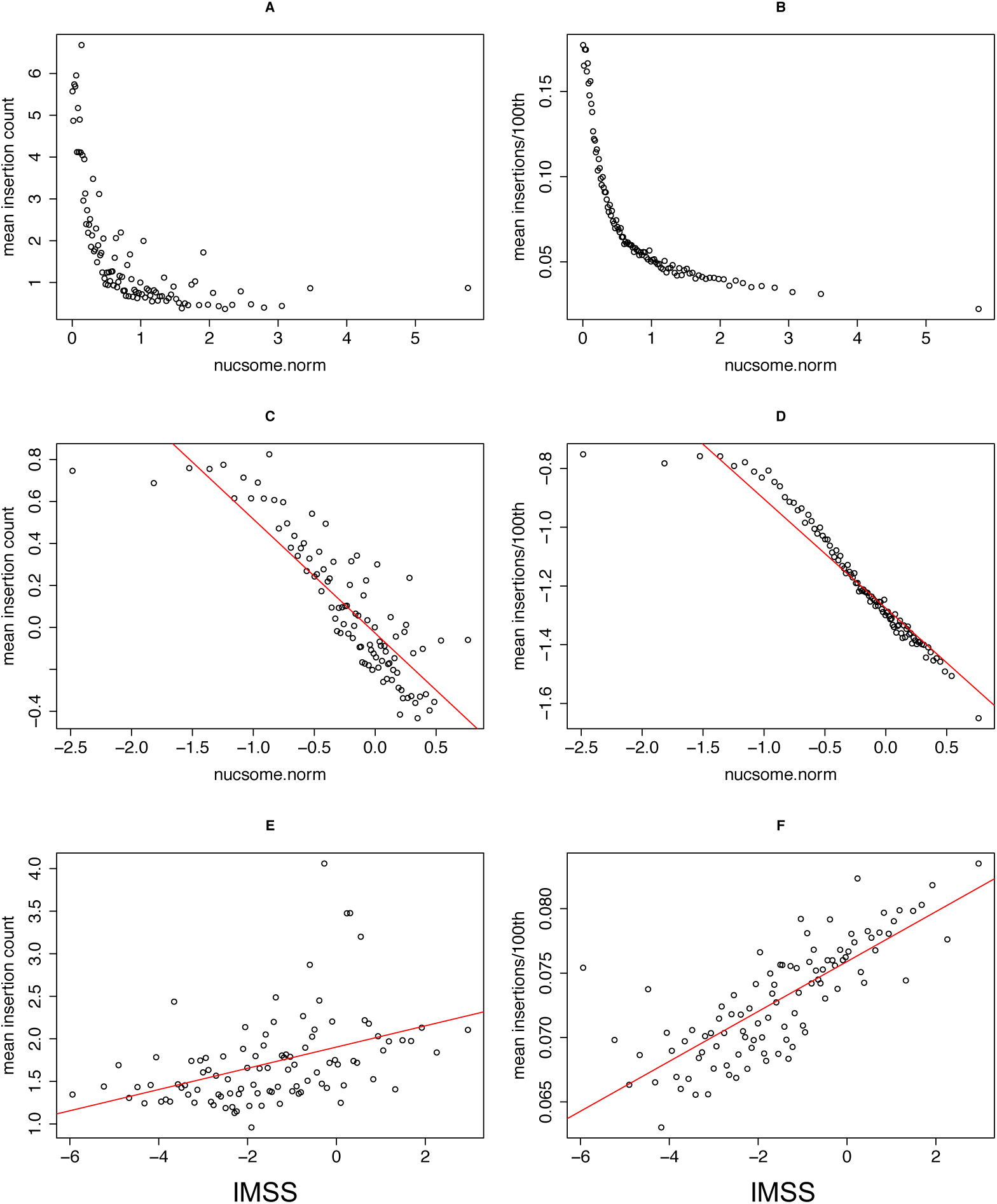
Relationships between insertion density, nucleosome density and the insertion motif similarity score. All plots show relationships with mean insertion count for sites with Hermes insertions (left panels) or mean insertions/site. In each case, the genome was divided into 100 partitions according to the measure on the *x* axis, and the insertion counts or insertion densities were calculated from these partitions. A) insertion counts plotted against normalised nucleosome density (nucsome.norm). B) insertion density plotted against normalised nucleosome density. C) log scale insertion counts plotted against log scale normalised nucleosome density. D) log scale insertion density plotted against log scale normalised nucleosome density. E) insertion counts plotted against insertion motif similarity score (IMSS). F) insertion density plotted against insertion motif similarity score.

**Supplementary fig. 6.**
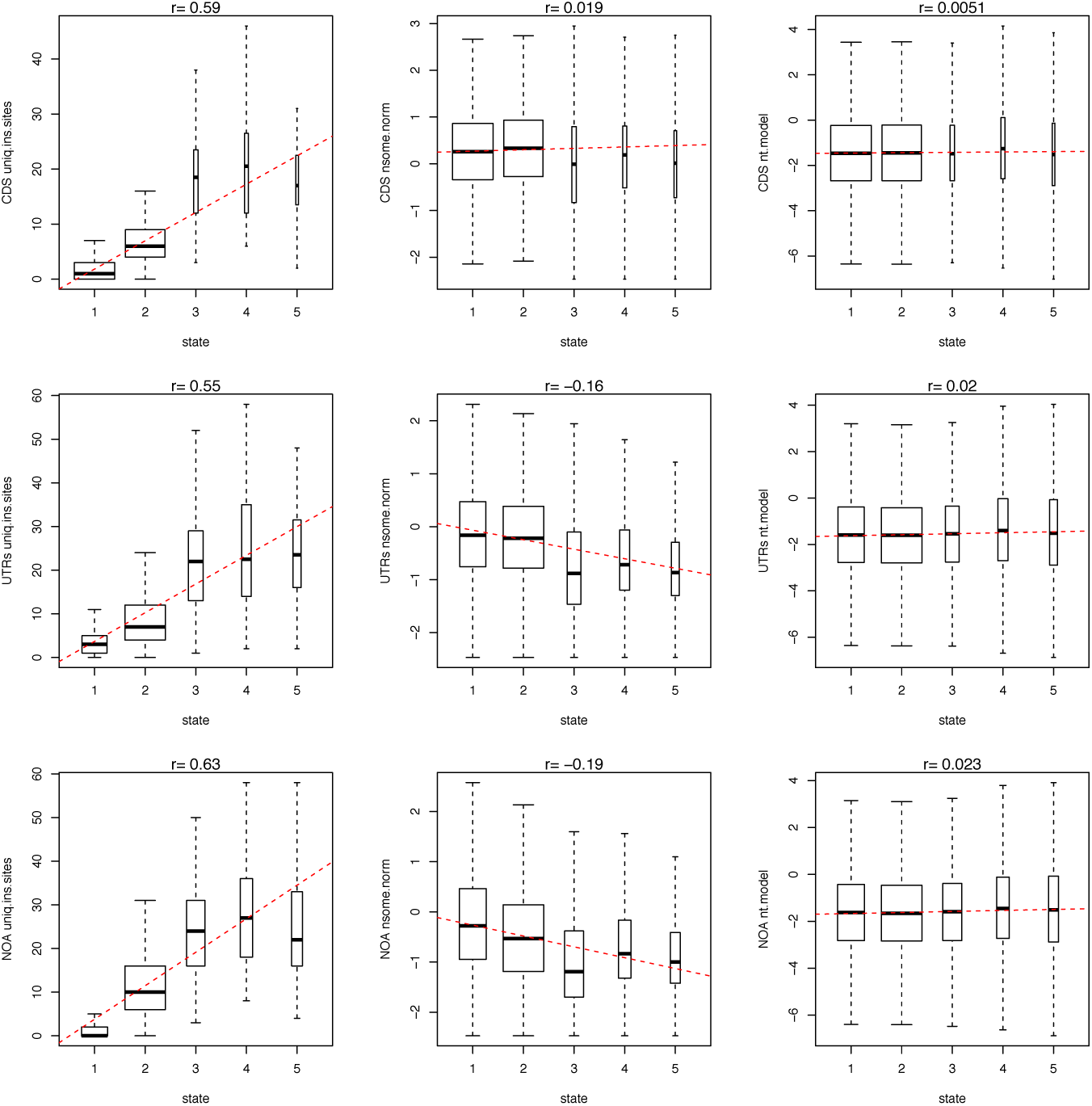
HMM states strongly depended on insertion density but only weakly correlated with nucleosome density and nucleotide motif. Top row; for coding regions we show the relationship between HMM states defined and insertion density (unique insertions/100 nt) (left panel), normalised nucleosome density (nsome.norm, middle panel) and the insertion motif similarity score (nt.model, right panel). Middle row; the same relationships for 5’ and 3’ untranslated regions. Lower row, the same relationships for regions with no annotations. In all cases Spearman rank correlations are shown above plots.

**Supplementary fig. 7.**
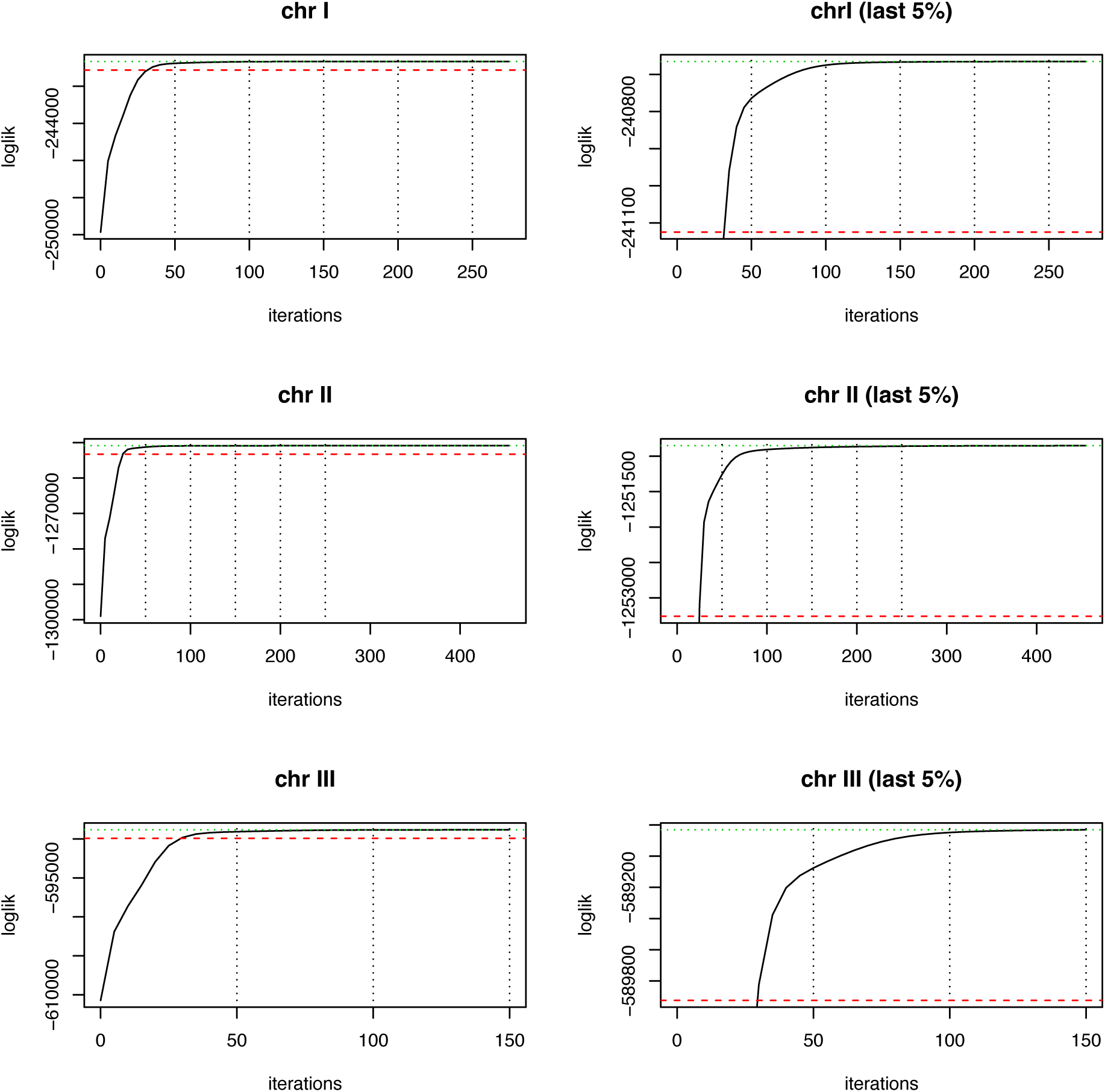
Log likelihoods for fits of HMM models improved little after 150 iterations. For sections of chromosomes I, II and III we show the log likelihood of the model fit to the data with successive iterations of the Viterbi algorithm. Left panels show the entire range of likelihoods, with red and green dashed lines showing the 95^th^ and 99^th^ percentiles. Right panels show the upper 5^th^ percentiles. Model fits improved little after 150 iterations.

**Supplementary fig. 8.**
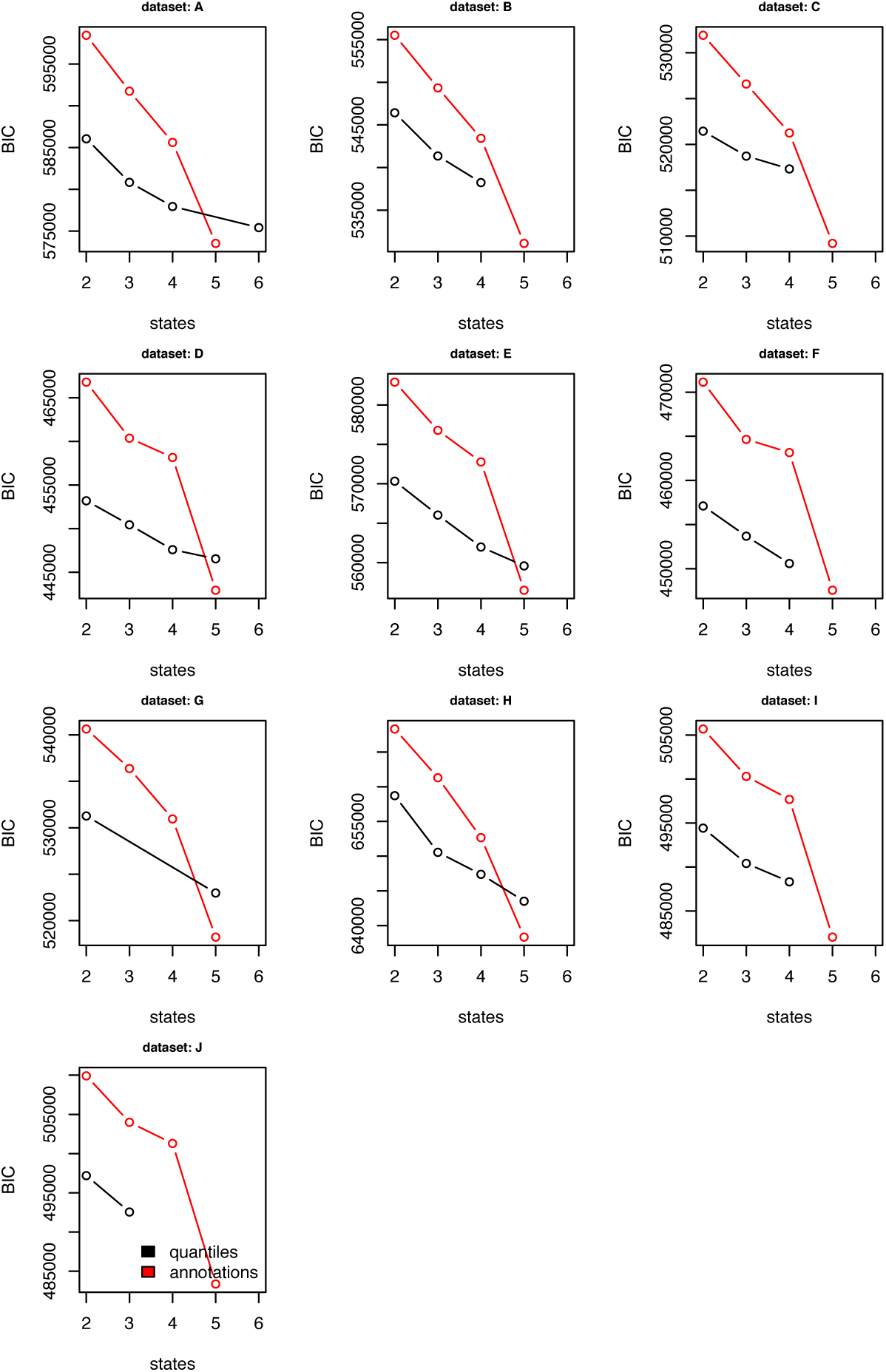
Bayesian information criterion scores (BIC) indicated that the 5-state annotation model was the best fit. For ten 100 kb fractions of the genome (data sets A – J), we show the BIC scores for model fitting with the depmixS4 package [42,43]. Red points show the annotation-based models from 2-5 states (see methods for state definitions). Black points show the quantile models, where training data is defined based on insertion density quantiles (unique insertions/100 nt). For example a three-state model used the first third of insertion-dense data to train S1, the second third to train S2, *etc*. The five-state model which was used for this analysis was trained on coding sequences of essential genes (S1), coding sequences of non-essential genes (S2), introns and untranslated regions (S3), and unannotated regions (S4), and sites with the highest 10% of unique insertions/100 nt (S5). The ten ‘test data’ subsets of the genome, each a 100 kb fraction as are follows: Chromosome I, 100001-200001, 1100001-1200001, 2100001-2200001, 3100001-3200001, Chromosome II, 100001-200001, 1100001-1200001, 2100001-2200001, 3100001-3200001 and Chromosome III, 100001-200001, 1100001-1200001 (test data sets A to J).

**Supplementary fig. 9.**
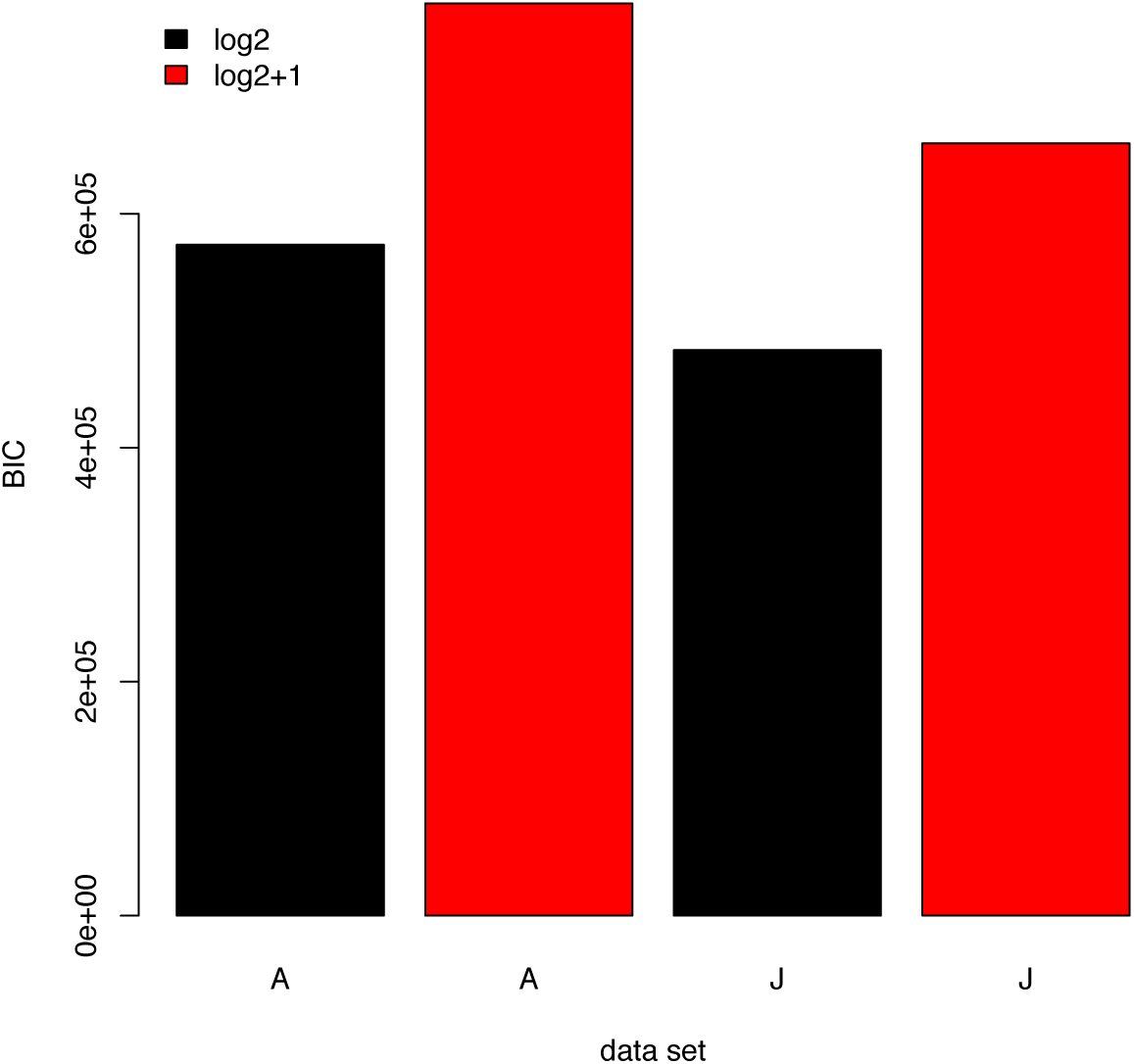
Excluding singleton insertions produced better model fits. HMM code used log_2_ of insertion counts (rounded to the nearest integer). Since log_2_(1) is zero, this treats sites with one insertion the same as sites with no insertions. Trails of the HMM code that used log_2_(insertions+1), where sites with 0 insertions have different value from those with 1, resulted in a worse fit to the model. For two of the test data sets (A, J), we show the BIC for models fitted with log_2_(insertions) and log_2_(insertions+1).

**Supplementary fig. 10.**
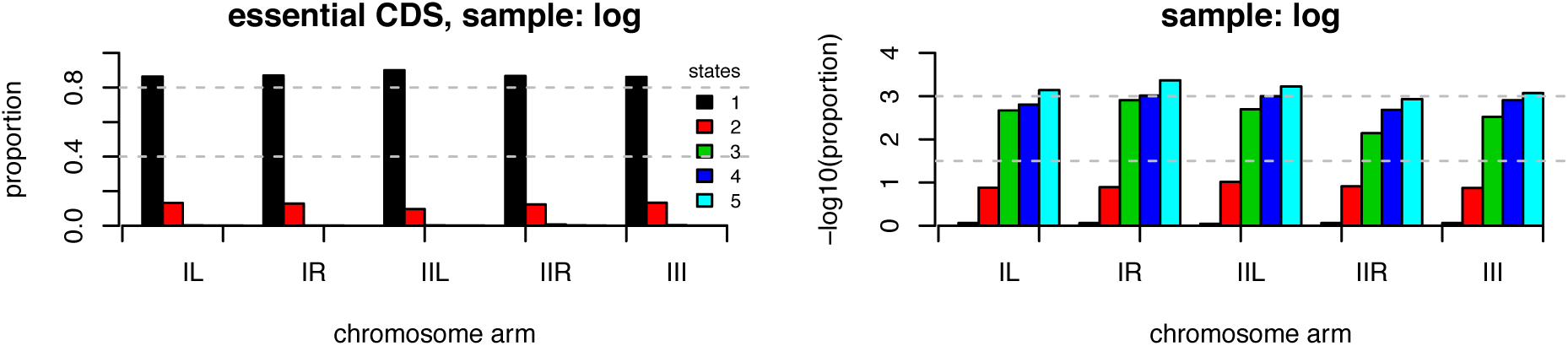
Separate fits to the model with different data resulted in similar distributions of states. Model fitting was performed on five subsets of the data; IL (left arm of chromosome I), IR (right arm of chromosome I), IIL (left arm of chromosome II), IIR (right arm of chromosome II), and III (all of chromosome III). The left panel shows the proportion of essential coding regions for each subset that were assigned to states 1-5, according to the key. Most were assigned to state 1 or 2. The right panel shows the –log10 of the proportion, which indicates that the less frequent states are also similarly distributed between subset model fits, supporting consistent convergence of the model between these genome subsets.

